# Interferon lambda 4 impacts broadly on hepatitis C virus diversity

**DOI:** 10.1101/305151

**Authors:** M Azim Ansari, Elihu Aranday-Cortes, Camilla LC Ip, Ana da Silva Filipe, Lau Siu Hin, Connor G G Bamford, David Bonsall, Amy Trebes, Paolo Piazza, Vattipally Sreenu, Vanessa M Cowton, STOP-HCV Consortium, Emma Hudson, Rory Bowden, Arvind H Patel, Graham R Foster, William L Irving, Kosh Agarwal, Emma C Thomson, Peter Simmonds, Paul Klenerman, Chris Holmes, Eleanor Barnes, Chris CA Spencer, John McLauchlan, Vincent Pedergnana

## Abstract

Type III interferons (IFN-λ) are part of the innate immune response to hepatitis C virus (HCV) infection however the specific role of IFN-λ4 and the nature of the viral adaption to this pressure have not been defined. Here we use paired genome-wide human and viral genetic data in 485 patients infected with HCV genotype 3a to explore the role of IFN-λ4 on HCV evolution during chronic infection. We show that genetic variations within the host *IFNL4* locus have a broad and systematic impact on HCV amino acid diversity. We also demonstrate that this impact is larger in patients producing a more active form of IFN-λ4 protein compared to the less active form. A similar observation was noted for viral load. We conclude that IFN-λ4 protein is a likely causal agent driving widespread HCV amino acid changes and associated with viral load and possibly other clinical and biological outcomes of HCV infection.

## Introduction

Hepatitis C virus (HCV) infects an estimated 71 million people worldwide^1^ and can lead to severe liver disease in chronically infected patients. The virus is highly variable and has been classified into 7 distinct genotypes, and further divided into 67 subtypes, based on nucleotide sequence identity^2^. The factors that have driven the evolutionary path of HCV are multifactorial but undoubtedly are also shaped by host genetics. Because of its major health burden, determining how both host and viral genetics contribute to the outcomes of infection is critical for understanding HCV-mediated pathogenesis^3^.

Using a systematic genome-to-genome approach in a cohort of chronically infected patients, we recently reported associations between an intronic single nucleotide polymorphism (SNP) rs12979860 in the interferon lambda 4 gene (CC vs. non-CC and herein referred to as *IFNL4* SNP or genotypes) and 11 amino acid polymorphisms on the HCV polyprotein^4^. This broad effects was unexpected since *IFNL4* is a member of the type III IFN family that act as cytokines as part of the innate immune system and therefore lack apparent epitope specificity^5^.

These associations between polymorphisms on the HCV polyprotein and host *IFNL4* genotypes are further intriguing given that variants within the *IFNL3/4* locus (including rs12979860 SNP) reportedly contribute to HCV clinical and biological outcomes, including spontaneous virus clearance, response to IFN-based treatment, viral load and liver disease progression^6–13^. It is possible that the associations between the outcomes of HCV infection and the *IFNL3/4* locus are inherently linked to its impact on the viral genome.

The intronic *IFNL4* SNP rs12979860 is in high linkage disequilibrium with other SNPs that may be more biologically relevant, including the exonic SNPs rs368234815 (r^2^=0.975 CEU population, 1000 Genomes dataset) in *IFNL4* and rs4803217 (r^2^=0.975 CEU population, 1000 Genomes dataset) in *IFNL3*. The SNP rs368234815 [ΔG>TT] causes a frameshift, abrogating production of IFN-λ4 protein^14^ and it is reported that the SNP rs4803217 [G>T] in 3’ UTR of *IFNL3* influences mRNA stability^15, 16^.

Moreover, an amino acid substitution (coded by the SNP rs117648444 [G>A]) in the IFN-λ4 protein, which substitutes proline for serine at position 70 (P70 and S70 respectively), reduces its antiviral activity *in vitro*^17^. Thus, the combination of SNPs rs368234815 and rs4803217 creates three haplotypes, one that does not produce IFN-λ4 protein (TT/G or TT/A; IFN-λ4-Null) and two that result in production of two IFN-λ4 protein variants (ΔG/G; IFN-λ4-P70 and ΔG/A; IFN-λ4-S70). Patients harbouring the impaired IFN-λ4-S70 variant display lower hepatic interferon-stimulated gene (ISGs) expression levels, which is associated with increased viral clearance following acute infection and a better response to IFN-based therapy, compared to patients carrying the more active IFN-λ4-P70 variant^18^.

In this study, we generated paired whole HCV genomes and genome-wide human SNP data from a cohort of 485 patients with self-reported white ancestry infected with HCV genotype (gt) 3a (411 from the BOSON^19^ cohort and 74 from the Expanded Access Programme (EAP) cohort^20^). We report that *IFNL4* genotypes have a widespread impact at polymorphic sites across the entire virus polyprotein. We also find an association with viral nucleotide content and certain dinucleotide frequencies, such as UpA. Finally, we demonstrate that IFN-λ4-S70 and IFN-λ4-P70 have different effect sizes on both viral load and viral amino acids. Together these observations suggest that IFN-λ4 is a major driver of HCV sequence diversity and clinical measures such as viral load.

## Results

### Viral principal components are associated with host *IFNL4* SNP

Paired human and viral genetic data were obtained for 485 HCV genotype 3a infected patients (N_BOSON_=411, N_EAP_=74, **Methods**). To control for both human and virus population structures, we performed principal component analysis (PCA) using host and viral genetic data separately (**Methods**). The host PCA defined a largely homogenous group corresponding to the self-reported white ancestry (**Supplementary Fig. 1a**). The first and second viral principal components (PCs) explained only 3% and 2% of HCV nucleotide diversity variance respectively (**Supplementary Fig. 1b**), as most of the observed evolution was on terminal branches of the phylogenetic tree (Fig. 1a). The viral sequences from the two cohorts 21 were non-randomly distributed on the tree^21^ and one clade was underrepresented in the EAP cohort sequences (Bayes factor = 249, **Methods** and **Supplementary Fig. 2a**). However, this observation was not reflected in host *IFNL4* genotypes, which was randomly distributed on the viral phylogenetic tree (Bayes factor = 1.1, **Supplementary Fig. 2b**).

**Figure 1:**
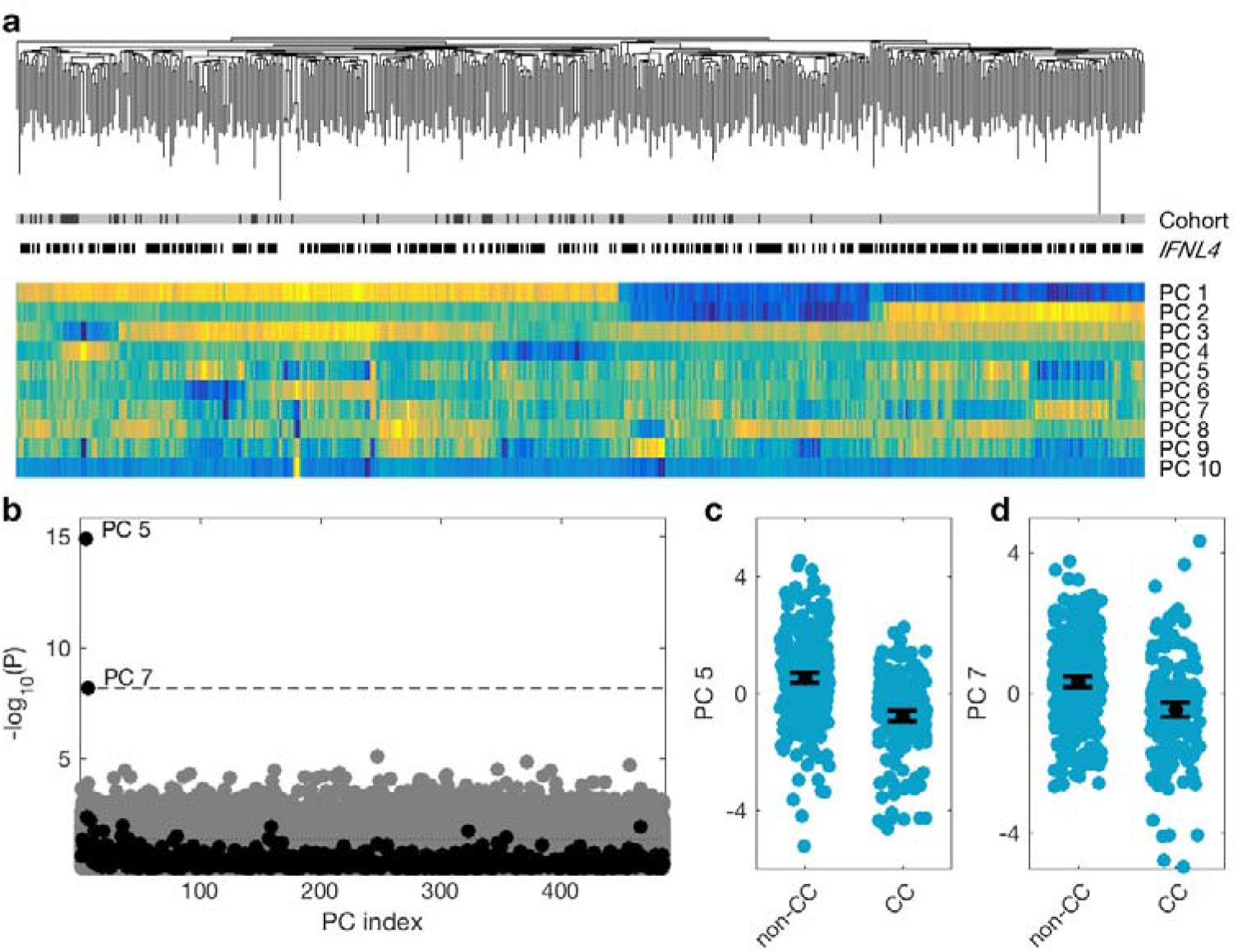
Association between viral PCs and *IFNL4* genotypes in the combined cohort. (**a**) Virus phylogenetic tree, cohort (black EAP, gray BOSON), *IFNL4* SNP (CC white, non-CC black) and the first 10 viral PCs (the colors are mapped such that dark blue represents the smallest number and bright yellow represents the largest number for each PC). (**b**) P-value of univariate association tests between viral PCs and the host SNPs. Black and gray dots are for association tests between the viral PCs and the *IFNL4* SNP and the 500 frequency-matched chosen SNPs respectively. Dashed line shows the 10% FDR line and the dotted line shows the nominal significance of *P*=0.05. Distribution of the fifth (**c**) and seventh (**d**) PCs stratified by the host *IFNL4* genotypes. Black dot and lines show the mean and its confidence interval for each group.

The first two viral PCs were clustered with clades on the virus phylogeny (Fig. 1a). The other PCs changed more gradually and were grouped to a lesser degree with specific clades on the tree. To explore any specific role for *IFNL4* SNP on viral diversity, we tested whether any viral PCs were associated with *IFNL4* genotypes in a univariate analysis and observed that 16 of the 485 viral PCs were nominally associated with *IFNL4* genotypes (*P*<0.05). At a 10% false discovery rate (FDR), the fifth and seventh PCs were associated with *IFNL4* genotypes (Fig. 1b-d) explaining 0.7% and 0.5% of the total variance in the nucleotide sequences (**Supplementary Fig. 1b**). The nucleotides of codon 2570 had the largest contribution (2.3%) to the fifth PC; this amino acid position was the most associated site with *IFNL4* SNP in our previous report (**Supplementary Fig. 3**). We then performed the same analysis in each cohort separately (**Supplementary notes**). In the BOSON cohort, we observed significant associations between three viral PCs and *IFNL4* genotypes (**Supplementary Fig. 4** and **5**). In the EAP cohort, none of the viral PCs were significantly associated with *IFNL4* genotypes, potentially due to small sample size. However, projecting the EAP viral sequences into the PCs axes of the BOSON cohort, we could predict *IFNL4* genotype in the EAP patients (area under the curve of 0.73). This indicated that the EAP viral PCs also carried information about the host genotypes (**Supplementary notes** and **Supplementary Fig. 6**).

To investigate whether the observed associations between *IFNL4* genotypes and viral PCs were due to unaccounted population structure, we selected 500 SNPs across the human genome with frequencies similar to the *IFNL4* SNP and tested for association between these SNPs and the viral PCs (Fig. 1b). If population structure was responsible for the observed association with *IFNL4* genotypes, we would expect these 500 SNPs to also correlate with viral PCs. At a 10% FDR, none of the viral PCs were associated with any of the 500 frequency-matched SNPs (performing 485×500 tests). Overall, these results indicate that the observed association between *IFNL4* genotypes and viral PCs is a consequence of biological interaction between the *IFNL3/4* locus and the virus sequences and not due to population structure or other systematic bias.

### *IFNL3/4* locus has a widespread impact on the viral polyprotein

A major advantage of determining entire HCV genomic sequence data is the possibility to perform footprinting analysis at a genome-wide scale. The nucleotide and amino acid frequencies at polymorphic viral sites in the two cohorts were similar and no systematic differences were observed (**Supplementary Fig. 7**). We tested the association between *IFNL4* genotypes and presence or absence of each amino acid at all variable sites (for amino acids present in at least 20 samples) on the HCV polyprotein, performing 977 tests at 471 sites. We observed that *P*-values were highly inflated (Fig. 2a) with a genomic inflation factor (λ) of 2.16. λ is defined as the ratio of the median of the empirically observed distribution of the association test statistic to the expected median, thus quantifying the extent of the bulk inflation in the observed statistic. Generally, an inflated λ value can reflect undetected sample duplications, unknown familial relationships, unaccounted and systematic technical bias and population structure but also potential enrichment in genuine associations.

**Figure 2:**
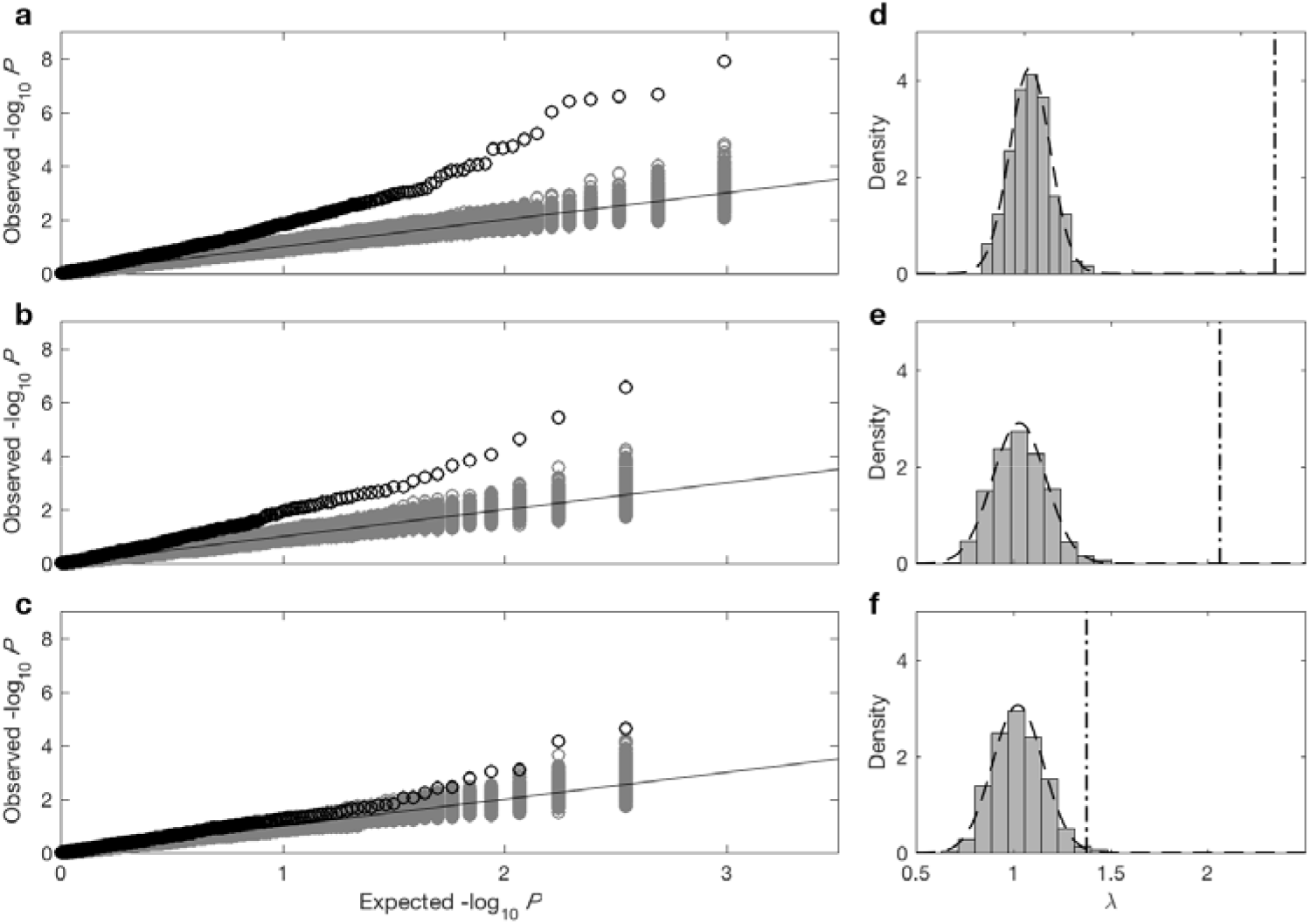
QQ-plots and genomic inflation factor (λ) distribution for association studies between viral amino acids and codons and *IFNL4* SNP rs12979860 and 500 SNPs chosen across the human genome frequency-matched to the *IFNL4* SNP. QQ-plots for association tests between the SNPs and viral amino acid (**a**) SNPs and change from the most common viral codon to (**b**) non-synonymous codons and (**c**) synonymous codons. First two viral PCs and first three host PCs were used as covariates in all three analyses. The black circles show the QQ-plot for the *IFNL4* SNP rs12979860 and the gray circles show the QQ-plot for the 500 frequency-matched SNPs. (**d, e** and **f**) Distribution of λs for association studies shown in **a, b** and **c** respectively. The dash dotted line indicates the λ for *IFNL4* SNP rs12979860 and the dashed line shows the normal distribution fitted to the λs in each analysis. Assuming the fitted normal is a reasonable estimate of the null distribution of λs, the P-value of observing λ of *IFNL4* SNP rs12979860 in each case is: **d)** 7.08×10^-34^, **e)** 2.65×10^-14^ and **f)** 3.43×10^-3^.

To ensure that the observed genomic inflation was not due to population structure or some other systematic bias, λ was estimated for the previously 500 selected SNPs (frequency-matched to *IFNL4* SNP) performing the same genomic association tests under three different models: without any covariates, including the first two viral PCs as covariates and including the first two viral and the first three host PCs as covariates. Inflation in the *P*-values was only observed for the *IFNL4* SNP and not the other 500 SNPs (Fig. 2a and **Supplementary Fig. 8**). Moreover including host and viral PCs as covariates had little impact on the results (**Supplementary Fig. 8**). We observed that the λ_*IFNL4*_ differed significantly from the distribution of λ under the null hypothesis of no association (estimated using λs from the 500 frequency-matched SNPs, *P*=7.08×10^-34^, Fig. 2d). In the BOSON cohort only analysis, similar results were observed (**Supplementary notes** and **Supplementary Fig. 9**).

Using the logistic regression association tests including two viral PCs and three host PCs, we found that 9% of variable sites (42/471) were associated with *IFNL4* SNP at 5% FDR, increasing to 16% of sites (76/471) at a 10% FDR (**Supplementary Fig. 10** and **Supplementary Table 1**). The most associated site was at position 2570 in the NS5B viral protein (*P*=1.32×10^-8^, log(OR)=1.19), as previously reported^4^. Notably, 26 of the 76 sites (34%) associated with the *IFNL4* SNP at a 10% FDR lie within the HCV E2 glycoprotein (**Supplementary notes** and **Supplementary Fig. 11**). However, we did not observe enrichment or depletion for association signals in any specific viral protein, or in previously reported HLA restricted epitopes in HCV genotype 3a^22^ (**Supplementary Table 2**). A meta-analysis of the independent analyses of the two cohorts showed similar results (**Supplementary notes** and **Supplementary Fig. 12**).

As association of *IFNL4* SNP was observed with both viral nucleotides (viral nucleotides PCA) and amino acids (viral amino acids GWAS), we explored nucleotide sequences at the codon level to distinguish the impact of the *IFNL4* SNP on viral nucleotides from its impact on viral amino acids. At each variable codon (where as well as the most common codon, there were at least 20 synonymous and 20 non-synonymous codons, N=348), we performed a logistic regression including two viral PCs and three host PCs to test for association between *IFNL4* SNP (and the 500 frequency-matched SNPs) and changes from the most common codon to synonymous and non-synonymous codons (**Methods**). We observed a highly significant inflation in the *P*-values of the association tests between the non-synonymous codon changes and *IFNL4* SNP (Fig. 2b and 2e, λ=2.06, *P*=2.65×10^-14^). The *P*-values for the association tests between the synonymous codon changes and the *IFNL4* SNP were slightly inflated (Fig. 2c and f, λ=1.38, *P*=3.43×10^-3^). This indicates that the observed association between *IFNL4* SNP and virus sequence diversity is most likely at the amino acid level, although a small impact on virus nucleotides cannot be excluded.

To further explore the viral nucleotide association, we estimated the dinucleotide frequencies for the different *IFNL4* genotypes (**Supplementary Fig. 13**). The UpA dinucleotide frequency (estimated as the ratio of observed to expected frequencies) was significantly lower in the *IFNL4* non-CC group compared to the CC group (*P*=1.5×10^-6^). By contrast, the UpG dinucleotide frequency was significantly higher in the *IFNL4* non-CC group compared to the CC group (*P*=1.5×10^-5^). The CpC and CpA dinucleotide frequencies were also significantly different between the *IFNL4* SNP groups using a Bonferroni correction for multiple testing (*P*<0.003). Similar results were observed by analyzing the cohorts independently (**Supplementary notes and Supplementary Fig. 14**)

### IFN-λ4 protein impacts viral amino acids and viral load

To refine the possible role of *IFNL4*, we explored the impact of the different haplotypes of the gene on HCV amino acid diversity and viral load. After imputing and phasing *IFNL4* SNPs rs368234815 and rs117648444, we observed three haplotypes: TT/G (IFN-λ4-Null); ΔG/G (IFN-λ4-P70) and ΔG/A (IFN-λ4-S70). HCV-infected patients were classified into three groups according to their predicted ability to produce IFN-λ4 protein: (i) no IFN-λ4 (two allelic copies of IFN-λ4-Null, N_BOSON_=145, N_EAP_=41), (ii) IFN-λ4-S70 (two copies of IFN-λ4-S70 or one copy of IFN-λ4-S70 and one copy of IFN-λ4-Null, N_BOSON_=48, N_EAP_=7), and (iii) IFN-λ4-P70 (at least one copy of IFN-λ4-P70, N_BOSON_=218, N_EAP_=26) (**Supplementary Table 3**).

Since IFN-λ4-S70 can be distinguished phenotypically from IFN-λ4-P70 both *in vivo* and *in vitro*, we examined whether the haplotypes had distinct effects on viral amino acid polymorphisms and clinical measures such as viral load. We estimated the effect size of IFN-λ4-S70 and IFN-λ4-P70 relative to the IFN-λ4-Null haplotype on the presence and absence of the 76 amino acids associated with *IFNL4* genotypes at 10% FDR. We found that the estimated effect sizes of IFN-λ4-S70 were consistently smaller than those for IFN-λ4-P70 (Fig. 3). Under the null hypothesis that there is no difference in the effect sizes of IFN-λ4-P70 and IFN-λ4-S70 alleles on viral amino acid polymorphisms, we would expect that the slope of the linear regression line (Fig. 3) to have a value of one. However, the estimated slope of the best-fit line was significantly different (slope = 0.77, *P*=9.6×10^-7^, Fig. 3).

**Figure 3:**
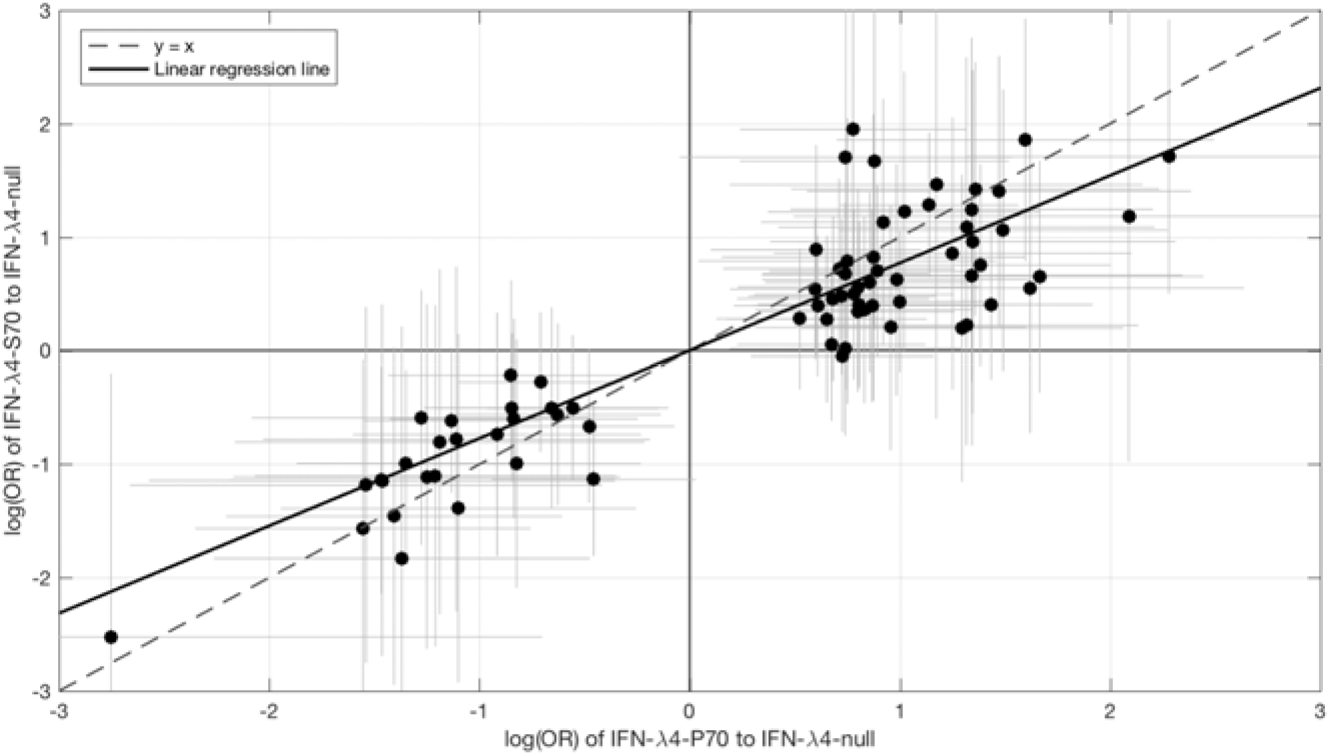
Comparing effect sizes (log(OR)) of host IFN-λ4 haplotypes (IFN-λ4-S70 and IFN-λ4-P70 relative to IFN-λ4-Null) on HCV amino acids. The circles shows the log(OR) estimates and the gray lines indicate the 95% confidence intervals. The dashed line is the y=x line which has a slope of one. The solid black line shows the linear regression line, which has a slope of 0.77 that is significantly different from one (y=x line, *P*=9.6×10^-7^).

We then investigated the effects of IFN-λ4 haplotypes on viral load. For this analysis, the EAP cohort was excluded as these patients had advanced liver disease with consistently lower viral loads relative to the BOSON cohort (**Supplementary Fig. 15**). We observed no difference in mean viral load between patients carrying IFN-λ4-S70 and IFN-λ4-Null haplotypes (*P*=0.61). However the viral load in patients carrying IFN-λ4-P70 was significantly lower than in the other two groups (*P*_*IFN-λ4-S70*_=1.6×10^-4^ and *P*_*IFN-λ4-Null*_=3.9×10^-10^), with IFN-λ4-P70 conferring an approximately 2.3-fold decrease in viral load compared to IFN-λ4-S70 (mean for IFN-λ4-P70=2, 905, 333, IFN-λ4-S70=6, 703, 875 and IFN-λ4-Null=6, 256, 523 IU/ml, Fig. 4a).

**Figure 4:**
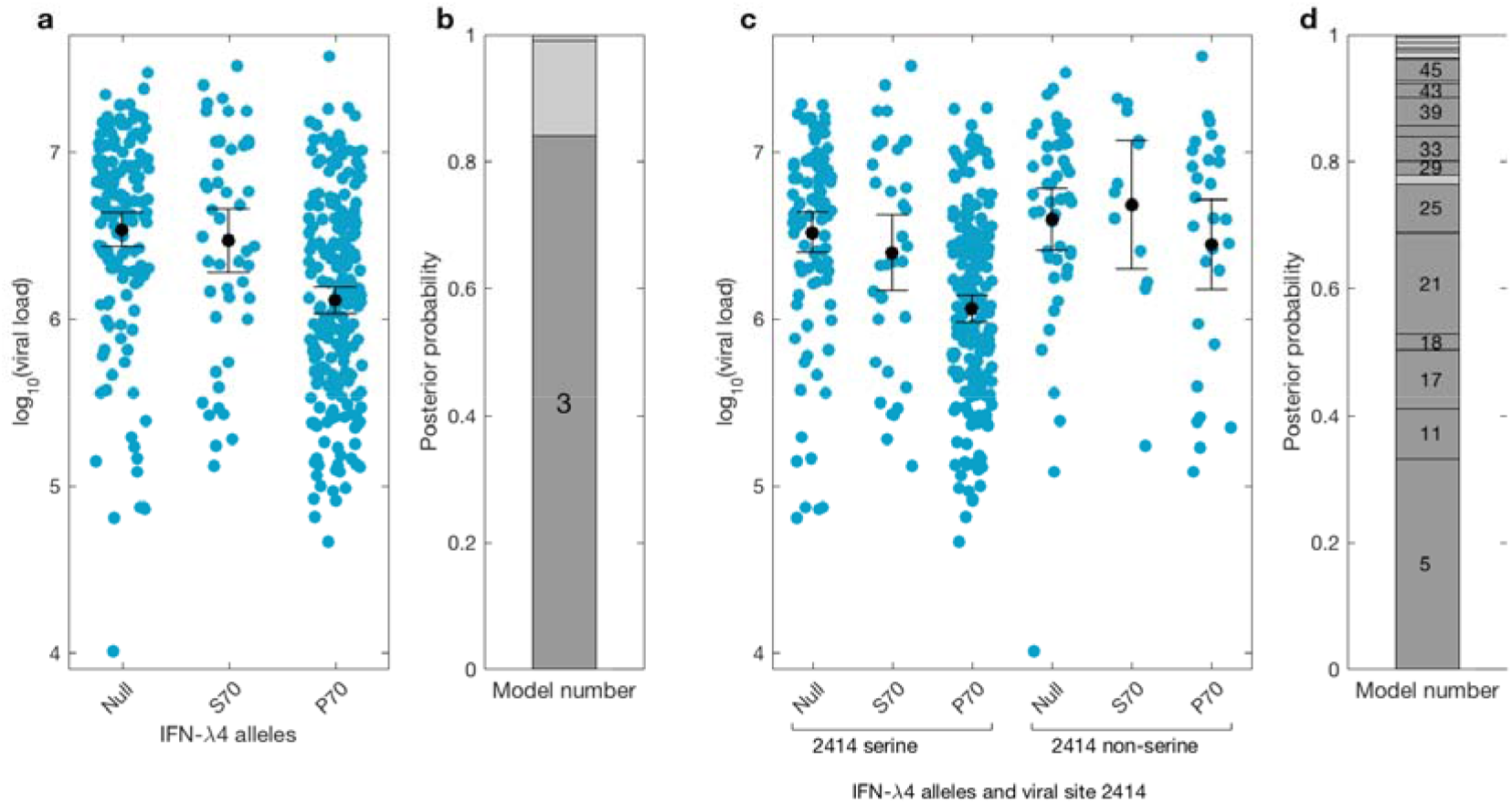
Bayesian model comparison of effect sizes of IFN-λ4 haplotypes on viral load. (**a**) Pretreatment viral load stratified by the host IFN-λ4 allele. The black dots and lines indicate the mean and confidence interval for each group. (**b**) The posterior probability of the five tested models from (**a**) stacked on top of each other. Models where the posterior probability is higher or lower than the prior probability are coloured as dark grey and light grey respectively. only model 3 has a posterior probability bigger than its prior probability and it assumes that the mean viral load is the same in IFN-λ4-Null and IFN-λ4-S70 groups while the mean viral load of IFN-λ4-P70 group is different from them. (**c**) Viral load stratified by the host IFN-λ4 allele and the presence and absence of serine in the viral amino acid site 2414. The black dots and lines indicate the mean and confidence interval for each group. (**d**) The posterior probability of the 58 tested models from (**c**) stacked on top of each other. Models where the posterior probability is higher or lower than the prior probability are coloured as dark grey and light grey respectively. Model 5 has the highest posterior probability and it assumes that the mean viral load is only different in “IFN-λ4-P70 + 2414 serine” group relative to the other groups.

We used a Bayesian approach to investigate the relationship between the effect sizes of the three IFN-λ4 haplotypes on viral load (Fig. 4b). In essence, this method weighs up the evidence that the genetic effects of the IFN-λ4-Null, IFN-λ4-S70 and IFN-λ4-P70 haplotypes are the same or not relative to each other (**Methods**). We tested five models; the effects of the three haplotypes are identical (model 1), the effects of IFN-λ4-P70 and IFN-λ4-Null are identical and different from the effect of IFN-λ4-S70 (model 2), the effects of IFN-λ4-S70 and IFN-λ4-Null are identical and different from the effect of IFN-λ4-P70 (model 3), all three haplotypes have different effect sizes (model 4) and the effects of IFN-λ4-P70 and IFN-λ4-S70 are the same but different from the effect of IFN-λ4-Null haplotype (model 5). Equal prior probabilities were used for all models. Model 3 had the highest posterior probability of 0.82 (Fig. 4b).

We had previously reported an interaction between *IFNL4* genotypes and the HCV amino acid at position 2414 in the NS5A protein associated with viral load^4^. HCV sequences were stratified by the viral amino acid at position 2414 (S2414), and host IFN-λ4 haplotypes. In viral sequences encoding serine, there was no significant difference in mean viral load between IFN-λ4-Null and IFN-λ4-S70 carriers (*P*=0.31, Fig. 4c), but both groups had a significantly higher viral load than IFN-λ4-P70 carriers (*P*_IFN-λ4-Null_=2.7×10^-9^; *P*_IFN-λ4-S70_=1.6×10^-10^,). However, no such association (*P*=0.49) between IFN-λ4 haplotypes and viral load was found in patients with a non-serine residue at this site (Fig. 4c) in agreement with our previous report^4^. We performed a Bayesian analysis that compared 58 possible models against each other (from all effect sizes being the same to all being different to each other). The model where only the “IFN-λ4-P70 + S2414” group had an effect size different from the other groups (model 5) had the highest posterior probability of 0.33 (Fig. 4b and **Supplementary notes**). Taken together, the combination of IFN-λ4-P70 and S2414 conferred a 2.6-fold decrease in viral load compared to IFN-λ4-S70 and S2414 (mean viral load for IFN-λ4-P70 and S2414=2, 376, 747 IU/ml, and mean viral load IFN-λ4-S70 and S2414=6, 093, 167 IU/ml).

## Discussion

Here, we show that genetic variants in the human *IFNL4* locus drive sequence change across the entire HCV polyprotein. We also report an association of the *IFNL3/4* locus with synonymous codon variants, suggesting that this locus might also affect the HCV genome at the nucleotide level. Finally, we report that IFN-λ4-S70 haplotype has a lesser impact on viral amino acid diversity and viral load compared to the more active IFN-λ4-P70 haplotype indicating that the *IFNL4* gene is likely not only a major driver of HCV amino acid variation but also modulates viral load in patients. Our findings extend the association between genetic variation in the *IFNL3/4* locus and outcome of HCV infection as well as hepatic disease^6–13, 18^.

We selected patients chronically infected with HCV genotype 3a and of self-reported white ancestry to limit the impact of human and viral population structures in our analyses. We observed significant associations between the fifth and seventh viral PCs (calculated using viral nucleotides) and host *IFNL4* SNP and that the *P*-values of association study between the host *IFNL4* SNP and the virus amino acids across the entire polyprotein were highly inflated. No such association or inflation was observed with 500 SNPs from across the human genome that were frequency-matched to the *IFNL4* SNP. This indicated that the observed impact of *IFNL4* SNP on the viral sequences was not due to population structure or other systematic bias. We conclude that *IFNL4* locus is an important driver of amino acid sequence change of HCV genotype 3a. Our studies provide a landmark for future analysis on whether *IFNL3/4* genetic variation also drives diversity in other HCV genotypes and subtypes across other ethnic populations.

To distinguish the effect of *IFNL4* SNP on viral amino acid from the nucleotide variability, we investigated the association of the *IFNL4* SNP with synonymous and non-synonymous codon changes. *IFNL4* SNP association tests with non-synonymous codon changes had highly inflated *P*-values, but we also observed a small inflation of *P*-values with synonymous codon changes. This indicated that although *IFNL4* SNP has a widespread impact at the amino acid level, it may also drive nucleotide diversity but to a lesser extent. To further explore the impact of the *IFNL4* variants on viral nucleotides, we investigated viral dinucleotide frequencies. The UpA dinucleotide frequency was significantly associated with the *IFNL4* genotypes; interestingly, ribonuclease L (RNase-L), an ISG that cleaves viral RNA to control viral infections in plants and animals^23^, targets both UpA and UpU dinucleotides^24^. Moreover, HCV genotype 1, which is relatively resistant to IFN-based therapy, has fewer UpA and UpU dinucleotides than the more IFN-sensitive HCV genotypes 2 and 3^25, 26^. However, we note that the *IFNL4* non-CC patients have a modest reduction (0.9%) in their viral UpA frequencies relative to the CC patients and that this reduction could be mediated by the widespread amino acid changes associated with the *IFNL4* SNP.

Due to high linkage disequilibrium in the *IFNL3/4* locus, it is difficult to distinguish the possible causal variant from correlated variants. We hypothesized that if *IFNL4* directs a bias in viral amino acid residues, then different effect sizes may be observed not only between IFN-λ4-P70 and IFN-λ4-Null but also IFN-λ4-S70, which produces a less active form of the protein. By imputing and phasing *IFNL4* SNPs in our cohort, we inferred the haplotypes consisting of the *IFNL4* SNPs rs368234815 (ΔG/TT) and rs117648444 (G/A). Using these data, we found that the IFN-λ4-S70 allele has a consistently smaller effect on viral amino acid variability relative to the IFN-λ4-P70 allele, which correlates with the reduced antiviral activity for IFN-λ4-S70 observed *in vitro*^17^. Moreover, the mean viral load in IFN-λ4-Null patients is similar to those carrying the IFN-λ4-S70 allele; by contrast, those carrying an IFN-λ4-P70 variant have a reduced viral load, which also correlates with *in vitro* data. Taken together, these observations reinforce the hypothesis that *IFNL4* is a functional gene with a major role in the HCV infection. We conclude that production of IFN-λ4 drives an altered immune response that mediates reduced viral load and increased impact on viral amino acid diversity.

In this study, we demonstrated by large-scale association studies that the *IFNL4* gene, a cytokine part of the innate immune response and therefore considered not to have specific effects at the amino acid level, can drive amino acid changes. We report that 4.2% (126/3021) of the HCV polyprotein amino acid are associated with *IFNL4* SNP (at a 20% FDR) and that the impact on amino acid variation is spread across the viral polyprotein. In comparison we previously reported that 5% of the HCV polyprotein was associated with *HLA* class I and II alleles^4^ (20% FDR) at the population level. The only other major driver of HCV amino acid variation is the B cell response, which is largely restricted to modifying epitopes on the envelope glycoproteins, in particular E2^27^. Thus, both arms of the adaptive immune system direct selection of amino acids encoded by HCV through pressure on epitopes recognized by T and B cell responses to infection.

Given that we did not observe any enhancement or depletion of association signals in a specific viral protein or in HLA restricted epitopes, we hypothesize that IFN-λ4 may exert its impact through a previously unknown mechanism or at more than one stage of the virus life cycle. We anticipate that this would result from distinct host responses in those who encoded variants that lead to IFN-λ4 synthesis as compared to individuals who carry the pseudogenized form of the gene. In common with other IFNs, IFN-λ4 induces a large number of ISGs, many of which are largely unstudied or poorly characterized. Since productive HCV infection in hepatocytes relies on a range of cellular pathways, it is likely that a spectrum of cellular functions are modified by genes stimulated by IFN-λ4. Within such an environment, it is possible that subtle selection of certain amino acids along the polyprotein will provide an advantage for viral entry, RNA replication, virion assembly and release. Further studies with appropriate *in vitro* models would address such questions and perhaps lead to identification of motifs in the mature viral proteins that contribute to the infection process.

Although there was no enrichment of associations comparing the structural with the non-structural proteins, the E2 glycoprotein contained the highest proportion of sites affected by *IFNL4* genotypes. From mapping these sites onto previously known functional domains on E2, we found that many residues mapped to either the hypervariable region 1 (HVR1) or the surface of the protein. Indeed, some sites coincided with epitopes that are targets for the antibody response or have a role in virus entry (**Supplementary Text** and **Supplementary Fig. 11**). Since the host response to HCV genotype 3a infection induces pathways including those affecting B cell development^28^, we cannot exclude the possibility that *IFNL4* genotypes either influence B cell response to infection or the process of virus binding and entry.

There are now multiple studies suggesting that *IFNL3/4* locus could be a key player in the defense against viruses other than HCV. In HIV-infected patients, the rs368234815 SNP has been associated with long-term non-progressor HIV-1 controllers^29^. In influenza virus infection, *IFNL3* SNP rs8099917 was associated with increased sero-conversion after influenza vaccination^30^. *IFNL4* variants have also been associated with bronchiolitis^31^, cytomegalovirus^32^ and Andes virus^33^ infections. These observations suggest that *IFNL4* possibly plays a role in many viral infections and immune related diseases in the liver and other organs. Investigating how IFN-λ4 (a cytokine without epitope specificity) drives amino acid selectivity in the HCV polyprotein would add a new dimension to how the human innate immune system interacts with viruses and controls infectious diseases.

## Online methods

### Patient cohorts

For this study we used patient data from the BOSON and EAP cohorts. To limit the potential impact of population structure, we restricted the analysis to patients of selfreported white ancestry infected with HCV genotype 3a for which we had obtained both host genome-wide SNP data and full-length HCV genome sequences. In total we have 485 patients in the study, 411 from the BOSON cohort and 74 from the EAP cohort.

Majority of the patients from the BOSON cohort have no or mild liver disease (compensated liver cirrhosis). The EAP cohort on the other hand consists of HCV-infected patients with advanced liver disease, the majority of whom had decompensated cirrhosis.

### Host genotyping and imputation

Informed consent for host genetic analysis was obtained from all patients. DNA samples from patients were genotyped using the Affymetrix UK Biobank array, as described elsewhere^4^. Phasing and imputation was performed using SHAPEIT2^34^ and IMPUTE2^35^ version 2.3.1 using default settings.

### Virus sequencing

RNA was isolated from 500 μl plasma using the NucliSENS magnetic extraction system (bioMerieux) and collected in 30 μl of kit elution buffer for storage at −80 °C in aliquots.

Libraries were prepared for Illumina sequencing using the NEBNext Ultra Directional RNA Library Prep Kit for Illumina (New England BioLabs) with 5 μl sample (maximum 10 ng total RNA) and previously published modifications of the manufacturer’s guidelines (v2.0)^36^, including fragmentation for 5 min at 94 °C, omission of actinomycin D at first-strand reverse transcription, library amplification for 18 PCR cycles using custom indexed primers^37^ and post-PCR clean-up with 0.85× volume Ampure XP (Beckman Coulter).

Libraries were quantified using Quant-iT PicoGreen dsDNA Assay Kit (Invitrogen) and analyzed using Agilent TapeStation with D1K High Sensitivity Kit (Agilent) for equimolar pooling; they were then re-normalized by qPCR using the KAPA SYBR FAST qPCR Kit (Kapa Biosystems) for sequencing. A 500-ng aliquot of the pooled library was enriched using the xGen Lockdown protocol from Integrated DNA Technologies (IDT) (Rapid Protocol for DNA Probe Hybridization and Target Capture Using an Illumina TruSeq or Ion Torrent Library (v1.0)) with equimolar-pooled 120-nt DNA oligonucleotide probes (IDT) followed by a 12-cycle, modified, on-bead, postenrichment PCR re-amplification step. The cleaned post-enrichment library was normalized with the aid of qPCR and sequenced with 151-base paired-end reads on a single run of the Illumina MiSeq using v2 chemistry.

De-multiplexed sequence-read pairs were trimmed of low-quality bases using QUASR (v7.0120)^38^ and of adaptor sequences using CutAdapt (version 1.7.1)^39^}, and they were subsequently discarded if either the read had less than 50 bases of remaining sequence or if both reads matched the human reference sequence using Bowtie (version 2.2.4)^40^. The remaining read pool was screened against a BLASTn database containing 165 HCV genomes^41^, which covered its diversity both to choose an appropriate reference and to select those reads that formed a population for de novo assembly with Vicuna (v1.3)^42^. The assembly was finished with V-FAT v1.0 (http://www.broadinstitute.org/scientific-community/science/projects/viral-genomics/v-fat). The population consensus sequence at each site was defined as the most common variant at that site among all of the patients.

### Statistical analysis

For the viral data, principle component analysis (PCA) was performed on the nucleotide data as follows. Tri- and quad-allelic sites were converted to binary variables. R (version 3.4.3, https://www.r-project.org) was used to perform the PCA using the prcomp function with default settings. Principle component analysis on the human genotype data was performed using flashpca^43^.

Whole-genome viral consensus sequences for each patient were aligned using MAFFT^44^ with default settings. This alignment was used to create a maximum-likelihood tree using RAxML^45^, assuming a general time-reversible model of nucleotide substitution under the gamma model of rate heterogeneity. The resulting tree was rooted at midpoint.

We used treeBreaker software^21^ (https://github.com/ansariazim/treeBreaker) to measure association between the virus phylogenetic tree and the host *IFNL4* SNP and the cohort ID. This software uses a Bayesian model to infer whether the phenotype of interest is randomly distributed on the tips of the tree and to estimate which branches have a distinct distribution of the phenotype of interest from the rest of the tree.

The univariate association between the *IFNL4* SNP (CC vs. non-CC) and the viral PCs was tested using logistic regression in R. We used the qvalue function from the qvalue package in R to perform the FDR analysis. As PCA was performed on the viral nucleotide sequences, to estimate the contribution of each codon to each PC, we added the contribution of all variables that were created for the nucleotides of that codon. The contribution of each variable to the PCs was estimated using function get_pca_var from the factoextra package in R.

To predict the host *IFNL4* genotypes in the EAP cohort from the viral PCs, we used the BOSON cohort as the training dataset and fit a logistic regression where the *IFNL4* genotypes was the response variable and the three viral PCs associated with it in the univariate analysis as the explanatory variables. Next we projected the EAP viral sequences into the same PC axis as the BOSON cohort analysis (using “predict” function in R). Next we used the projected EAP PCs and the estimated model parameters from the BOSON cohort to predict the host *IFNL4* genotypes (using “predict” function in R). Finally we used “ROCR” package in R to compare the predicated *IFNL4* genotypes to the actual genotypes in the EAP cohort and to calculate the area under the curve for the classifier.

To choose 500 SNPs across the human genome with similar frequency as the *IFNL4* SNP rs12979860, we used Fisher’s exact test to compare all SNPs against the *IFNL4* SNP (2×3 contingency table where the columns indicate the frequencies of 1, 2 and 3 copies of the minor allele and the rows are the *IFNL4* SNP and the target SNP counts) and chose the 500 SNPs with the largest p-values (least significance). SNPs in the *IFNL3-IFNL4* region were not included.

To perform the association tests between the virus amino acids and the host SNPs we used logistic regression in R. We investigated presence and absence of each amino acid at all variable sites, given that the amino acid was present in at least 20 samples. The presence and absence of the viral amino acid was used as the response variable and the host SNP coded as 0 and 1 based on presence and absence of minor allele as the explanatory variable (the same coding as the *IFNL4* SNP CC vs. non-CC). When the analysis included host and viral PCs, they were included as explanatory variables in the logistic regression. The genomic inflation factor (λ) was calculated as the median of the observed chi-squared test statistics divided by the median of the chi-squared distribution with one degree of freedom.

To test for enrichment or depletion of the association signals in a viral protein or the epitope regions, we used Fisher’s exact test. Each tested site is either within the target region or not and it is either classified as significant or not. The resulting 2×2 contingency table was tested using fisher.test function in R.

To perform meta-analysis of association between *IFNL4* SNP and the viral amino acids, the BOSON and EAP cohorts were analysed independently using logistic regression as previously described. We used fixed effects method to perform metaanalysis. To analyse how often the effect sizes are consistent between the two cohorts (**Supplementary notes**), we used a binomial test. Under the null hypothesis that the effects identified in the BOSON cohort are false positives, we would have expected the direction of the effect in the two cohorts to be the same 50% of the time. The sites were sorted in increasing P-value order from the BOSON cohort. For each site, we used the most associated amino acid in the BOSON cohort as the target amino acid to get the direction of effect. Increasing the number of sites one at a time, we counted the number of times that the direction of effect sizes were consistent in the two cohorts and the total number of sites being analysed and used the binomial test with probability of success of 0.5.

To separate the impact of the *IFNL4* SNP on amino acids from the nucleotides we investigated the nucleotide sequences at the codon level. At each codon (where there were at least 20 synonymous and 20 non-synonymous codons for the most common codon) we used logistic regression to test for association between *IFNL4* SNP (CC vs. non-CC) and the changes from the most common codon to synonymous and non-synonymous codons. The *IFNL4* SNP was the response variable and the codons were used as a categorical explanatory variable with three levels. The effect sizes (log(OR)) were estimated for the synonymous and non-synonymous codons relative to the most common codon. We used two viral PCs and three host PCs as covariates in this analysis.

To calculate the dinucleotide frequencies, the observed proportion of each dinucleotide was normalized by its expected proportion (assuming the nucleotides are independent the expected proportion can be calculated by multiplying the observed proportions for the relevant nucleotides). To test for association with the *IFNL4* genotypes we used a linear regression were the normalized dinucleotide proportions were used as the response variable and the *IFNL4* genotype as a categorical explanatory variable. We used two viral PCs and three host PCs as covariates.

To estimate the effect of the IFN-λ4 protein variants on the HCV amino acids, we used the 76 sites that were associated with *IFNL4* SNP at 10% FDR. Each patient was categorised to one of the three groups; not producing IFN-λ4 (IFN-λ4-Null), producing IFN-λ4-P70 and producing IFN-λ4-S70. We then used logistic regression to estimate the effect sizes (log(OR)) for IFN-λ4-P70 and IFN-λ4-S70 on the virus amino acids relative to the IFN-λ4-Null. The presence and absence of the reported viral amino acid was used as the response variable and the host IFN-λ4 status was used as the explanatory variable with the IFN-λ4-Null used as the base level and the log odds ratios for the IFN-λ4-P70 and IFN-λ4-S70 were estimated relative to the IFN-λ4-Null base level. We included two viral PCs and three host PCs as cofactors to account for possible population structure. To test whether the effect sizes of IFN-λ4 variants on viral amino acids are the same, we used the above estimated effect sizes and fit a linear regression line to it. One viral site was excluded from this analysis as it had unreliable effect size estimate (log(OR) = −17) for the IFN-λ4-S70 variant. Under the null hypothesis that the IFN-λ4-P70 and IFN-λ4-S70 have the same effect sizes, we would expect that the linear regression line to have a slope of one. To test whether the slope of the fitted line is different from one, we used R to fit a linear regression line that goes through the origin and used the offset function (F-test).

To assess the evidence for whether the mean viral load is different in the three patient groups of IFN-λ4-Null, IFN-λ4-P70 and IFN-λ4-S70, we used a Bayesian framework to perform model comparison (see **Supplementary notes** for further details). The models we considered comprised fixed and independent effects between the IFN-λ4 variants. We standardised the log10(viral load) so that it had a mean of zero and standard deviation of one. We used linear regression to get maximum likelihood estimates of the effects of IFN-λ4-S70 and IFN-λ4-P70 variants relative to the IFN-λ4-Null variant. The estimates were adjusted for cirrhosis status and population structure (including two viral PCs and three host PCs in the regression as covariates). For each effect size we assumed a normally distributed prior on the log(OR) of association with mean of zero. The prior covariance matrix determines the prior model assumptions. The elements of the covariance matrix were chosen such that the relevant prior model is set (see **Supplementary notes** for details).

To assess the evidence for interaction between host IFN-λ4 variants and viral amino acid site 2414, we used the same Bayesian framework detailed above. The patients were grouped into six categories based on the host IFN-λ4 variants and the presence or absence of serine at viral site 2414. We standardised the log10(viral load) so that it had a mean of zero and standard deviation of one. We used linear regression to get maximum likelihood estimates of the effects of “IFN-λ4-Null + 2414 not serine”, “IFN-λ4-P70 + 2414 not serine”, “IFN-λ4-P70 + 2414 serine”, “IFN-λ4-S70 + 2414 not serine”, “IFN-λ4-S70 + 2414 serine” groups relative to the “IFN-λ4-Null + 2414 serine” group. The estimates were adjusted for cirrhosis status and population structure (including two viral PCs and three host PCs in the regression as covariates). The prior covariance matrix determines the prior model assumptions. The elements of the covariance matrix were chosen such that the relevant prior model is set (see **Supplementary notes** for details).

## Materials & Correspondence

Correspondence and material requests should be addressed by contacting STOP-HCV http://www.stop-hcv.ox.ac.uk/contact.

## Author contributions

M.A.A and E.A.C contributed equally; J.M and V.P jointly supervised research; M.A.A, E.A.C, C.C.A.S, J.M and V.P conceived and designed the experiments; M.A.A, E.A.C, A.S.F, L.S.H, C.I, D.B, A.T, P.P, V.S and V.P performed the experiments; M.A.A, C.C.A.S and V.P performed statistical analysis; M.A.A, E.A.C, V.M.C, A.H.P, C.C.A.S, J.M and V.P analysed the data; M.A.A, E.A.C, A.S.F, L.S.H, C.G.G.B, C.I, D.B, A.T, P.P, V.S, V.M.C, E.M, R.B, P.K, A.H.P, G.R.F, W.L.I, K.H, P.S, E.T, E.B contributed reagents/materials/analysis tools; M.A.A, E.A.C, E.B, C.C.A.S, J.M and V.P wrote the paper.

## Acknowledgements

The authors would like to thank Gilead Sciences for the provision of samples and data from the BOSON clinical study for use in these analyses and HCV Research UK (funded by the Medical Research Foundation [C0365]) for their assistance in handling and coordinating the release of samples for these analyses. The authors would also like to thank Daniel J Wilson and Jacques Fellay for helpful comments.

This work was funded by a grant from the Medical Research Council (MR/K01532X/1 – STOP-HCV Consortium). The work was supported by Core funding to the Wellcome Trust Centre for Human Genetics provided by the Wellcome Trust (090532/Z/09/Z). E.C.T is funded by Wellcome Trust as a clinical fellow (102789/Z/13/Z). E.B is funded by the MRC as an MRC Senior Clinical Fellow with additional support from the Oxford NHIR BRC as a principle fellow. Professor Barnes is a National Institute for Health Research (NIHR) Senior Investigator. P.K is funded by the Oxford Martin School, NIHR Biomedical Research Centre, Oxford, by the Wellcome Trust (109965MA) and NIH (U19AI082630). C.C.A.S is funded by the Wellcome Trust (097364/Z/11/Z). Work in AHP and JM’s laboratories are supported by the MRC Core funding (MC_UU 12014/2). The views expressed in this article are those of the author(s) and not necessarily those of the NHS, the NIHR, or the Department of Health.

